# Correlation of mitochondrial TOM core complex *stop-and-go* and open-closed channel dynamics

**DOI:** 10.1101/2021.09.28.462098

**Authors:** Shuo Wang, Lukas Findeisen, Sebastian Leptihn, Mark I. Wallace, Marcel Hörning, Stephan Nussberger

## Abstract

The role of lateral diffusion of proteins in the membrane in the context of function has not been examined extensively. Here, we explore the relationship between protein lateral diffusion and channel activity of the general protein import pore of mitochondria (TOM-CC). Optical ion flux sensing through single TOM-CC molecules shows that TOM-CC can occupy three ion permeability states. Whereas freely diffusing TOM-CC molecules are preferentially found in a high permeability state, physical tethering to an agarose support causes the channels to transition to intermediate and low permeability states. This data shows that combinatorial opening and closing of the two pores of TOM-CC correlates with lateral protein diffusion in the membrane plane, and that the complex has mechanosensitive-like properties. This is the first demonstration of β-barrel protein mechanosensitivity, and has direct conceptual consequences for the understanding of the process of mitochondrial protein import. Our approach provides a novel tool to simultaneously study the interplay of membrane protein diffusion and channel dynamics.

## Introduction

The TOM complex of the outer membrane of mitochondria is the main entry gate for nuclear-encoded proteins from the cytosol into mitochondria (Wiedemann and Pfanner, 2017). It does not act as an independent entity, but in a network of interacting protein complexes, which transiently cluster in mitochondrial outer- and inner-membrane contact sites (Pfanner et al., 2019; Scorrano et al., 2019; Wurm et al., 2011). Recent studies suggest a functional crosstalk between TOM and chaperones, which guide preproteins from the cytosol to mitochondria (Becker et al., 2019; Namba, 2019), cytosolic ribosomes (Gold et al., 2017), and the endoplasmic reticulum (Becker et al., 2019; Namba, 2019). For proteins destined for integration into the lipid bilayer of the inner mitochondrial membrane, TOM transiently cooperates with components of the inner membrane protein translocase TIM22. Proteins en route to the mitochondrial matrix require supercomplex formation with the inner membrane protein translocase TIM23 (Chacinska et al., 2005; Gold et al., 2014; Mokranjac et al., 2009). Thus, depending on the activity of mitochondria, the lateral mobility of TOM within the mitochondrial outer membrane may be fundamental to the different import needs of the organelle (Pfanner et al., 2019). The relationship between the mode of lateral mobility (Jacobson et al., 2019) and TOM protein function, however, has not yet been explored. In particular, it is important to know whether the main entry gate for nuclear-encoded proteins is open or closed when associated with structures at the membrane periphery, as this could have direct conceptual implications for mitochondrial protein import.

Detailed insights into the molecular architectures of the TOM core complex (TOM-CC) from *N. crassa* (Bausewein et al., 2017), *S. cerevisiae* (Araiso et al., 2019; Tucker and Park, 2019) and human (Wang et al., 2020) mitochondria have been obtained by cryo-electron microscopy (cryoEM). All three structures show well-conserved symmetrical dimers, where the monomer comprises five membrane protein subunits. Each of the two transmembrane β-barrel domains of the protein-conducting subunit Tom40 interacts with one subunit of Tom5, Tom6, and Tom7, respectively. Two central transmembrane Tom22 receptor proteins, reaching out into the cytosol and the mitochondrial inter membrane space (IMS), connect the two Tom40 pores at the dimer interface. The path of polypeptides through theses pores is considered identical to the path of ions through the complex.

Previous studies reported that the TOM complex channels from wild-type yeast mitochondria are mainly in closed state (van Wilpe et al., 1999). However, TOM isolated from mutant *tom22Δ* mitochondria without Tom22 was mainly found in open state (van Wilpe et al., 1999), resembling that observed with purified Tom40 (Ahting et al., 2001; Hill et al., 1998). This indicated that the “multifunctional organizer” Tom22 of the mitochondrial protein translocase negatively regulates the open probability of the Tom40 channels of the TOM-CC (van Wilpe et al., 1999). Nevertheless, the molecular mechanism of how Tom22 affects the open-closed channel activity of the TOM machinery has remained an open question.

Most studies that reported on the dynamic and functional properties of the TOM-CC channel have been based on ion current measurements through single TOM-CC channels in planar lipid membranes under application of a membrane potential (Δψ_m_) (Ahting et al., 1999; Becker et al., 2005; Hill et al., 1998; Künkele et al., 1998; Kuszak et al., 2015; Mager et al., 2011; Mahendran et al., 2013). TOM-CC has been found to switch between a complex set of conductance states under this experimental setting. Tom22 has been proposed to contribute additional flexibility to the complex by reducing the energy required for transitions between these states (Poynor et al., 2008; Romero-Ruiz et al., 2010). However, the physiological significance of these voltage-dependent transitions has been controversially discussed because the critical voltage above which TOM-CC channels close is significantly larger than any possible Donnan potential (Δψ_Don_) across the outer mitochondrial membrane (Benz et al., 1990; Lemeshko and Lemeshko, 2000; Lemeshko, 2002).

In this work, we simultaneously probe and correlate the lateral mobility and the ion flux though single TOM-CC molecules by means of non-invasive Electrode-free Optical Single Channel Recording (Ef-oSCR) (Demuro and Parker, 2005; Huang et al., 2015; Leptihn et al., 2013; Wang et al., 2018; Yuqin Wang et al., 2019). Possible changes in ion flux associated with conformational changes of the TOM-CC are only caused by thermal fluctuations and the nature of lateral diffusion of the TOM-CC in the membrane. This leads to new insights into the open-closed dynamics of the TOM-CC channel and the corresponding nature of lateral movement, which are difficult to illuminate by voltage-clamp, biochemical assays or structural analysis (Davis et al., 2021).

We find that freely diffusing TOM-CC molecules stall when interacting with structures adjacent to the membrane, ostensibly due to interaction between the extended polar domains of Tom22 and a supporting agarose layer. Concomitantly, with transient or permanent suspension of movement, the TOM-CC switches from active (both pores open) to weakly active (one pore open) and inactive (both pores closed) states.

The strong temporal correlation between lateral mobility and channel activity suggests that TOM-CC function is highly sensitive to lateral diffusion and mechanical constraints. Taken alongside recent cryo-electron microscopy of this enigmatic complex, we argue that these dynamics provide a new functionality of the TOM-CC that has not been anticipated before. From a general perspective, for the best of our knowledge this is the first demonstration of β-barrel protein mechanoregulation and the causal effect of lateral diffusion. So far, mechanosensitivity has only been observed in primarily α-helical protein channels. The applied experimental approach can be easily transferred to other systems, where channel activity additionally to lateral membrane diffusion is of importance.

## Results

### Visualizing the open-closed channel activity of TOM-CC

Owing to the unresolved complexity of the mitochondrial outer membrane, TOM-CC was isolated from a *N. crassa* strain, that carries a version of subunit Tom22 with a hexahistidine tag at the C-terminus (Fig. 1A), and reconstituted into a well-defined supported lipid membrane (Kiessling et al., 2015; Sackmann, 1996; Tanaka and Sackmann, 2005). Droplet interface membranes (DIBs) were created through contact of lipid monolayer-coated aqueous droplets in a lipid/oil phase and a lipid monolayer on top of an agarose hydrogel (Huang et al., 2015; Leptihn et al., 2013; Wang et al., 2018; Yuqin Wang et al., 2019). The *cis* side of the membrane contained Ca^2+^-ions, while having a Ca^2+^-sensitive fluorescent dye (Fluo-8) at the *trans* side. Ca^2+^-ion flux through individual TOM-CCs was measured by monitoring Fluo-8 emission in close proximity to the membrane using TIRF microscopy in the absence of membrane potential (Ef-oSCR) to avoid voltage-dependent TOM-CC gating (Figs. 1B and 1C).

**Fig. 1.**
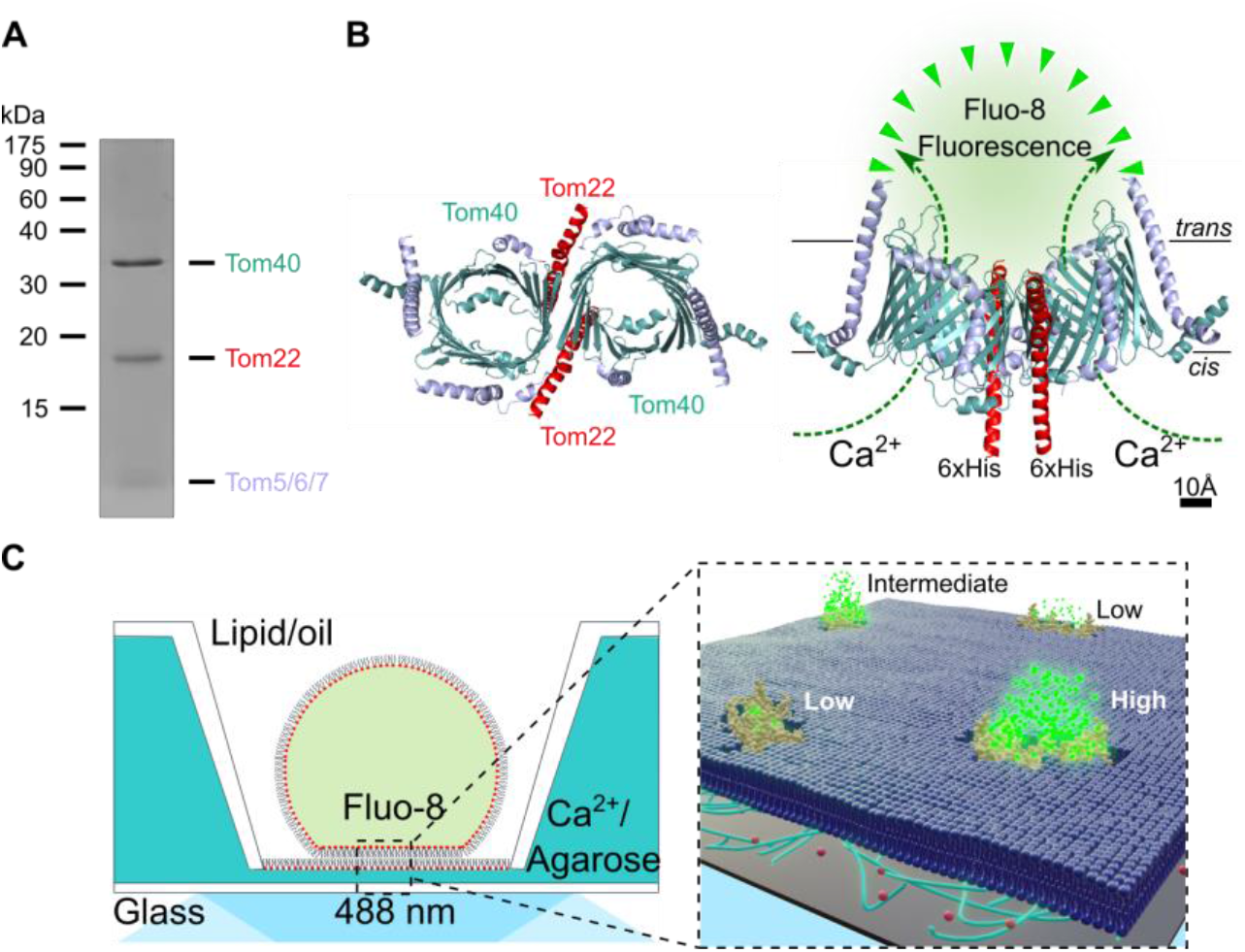
Scheme for tracking single TOM-CC molecules and imaging their ion channel activity. (A) TOM-CC was isolated from mitochondria of a *Neurospora* strain carrying a Tom22 with a hexahistidinyl tag (6xHis). Analysis of purified protein by SDS-polyacrylamide gel electrophoresis (SDS-PAGE) followed by Coomassie Blue staining revealed all known subunits of the core complex, Tom40, Tom22, Tom7, Tom6 and Tom5. The small subunits Tom7, Tom6 and Tom5 are not separated by SDS-PAGE. (B) Atomic model based on the cryoEM map of *N. crassa* TOM core complex (EMDB, EMD-3761; (Bausewein et al., 2020, 2017)). The ionic pathway through the two aqueous β-barrel Tom40 pores is used to optically study the open-closed channel activity of individual TOM-CCs. Left, cytosolic view; right, side view; *cis*, mitochondria intermembrane space; *trans*, cytosol. Tom7, Tom6 and Tom5 are not labeled for clarity. (C) Single molecule tracking and channel activity sensing of TOM-CC in DIB membranes using electrode-free optical single-channel recording (Ef-oSCR). Left: Membranes are created through contact of lipid monolayer-coated aqueous droplets in a lipid/oil phase and a lipid monolayer on top of an agarose hydrogel. The *cis* side of the membrane contained Ca^2+^-ions, while having a Ca^2+^-sensitive fluorescent dye (Fluo-8) at the *trans* side. Right: Ca^2+^-ion flux through individual TOM-CCs from *cis* to *trans* is driven by a Ca^2+^ concentration gradient, established around the two Tom40 pores, and measured by monitoring Fluo-8 emission in close proximity to the membrane using TIRF microscopy. Fluorescence signals reveal the local position of individual TOM-CCs, which is used to determine their mode of lateral diffusion in the membrane. The level of the fluorescence (high, intermediate and low intensity) correlates with corresponding permeability states of a TOM-CC molecule. A 100x TIRF objective is used both for illumination and imaging. Green dots, fluorescent Fluo-8; red dots, Ca^2+^ ions.

Contrary to the classical single molecule tracking approach using single fluorescently labeled proteins, we have an almost instantaneous update of fluorophores close to the TOM-CC nanopores. This enables a spatiotemporal tracking of individual molecules with much higher accuracy and longer observation time up to a couple of minutes.

Upon TIRF illumination of membranes with 488 nm laser light, single TOM-CC molecules appeared as high-contrast fluorescent spots on a dark background (Fig. 2A). High (S_H_), intermediate (S_I_) and low (S_L_) intensity levels were indicating Ca^2+^-flux through the TOM-CC in three distinct permeability states. No high-contrast fluorescent spots were observed in membranes without TOM-CC. The fact that the TOM-CC is a dimer with two identical β-barrel pores (Fig. 1B) suggests that the high and intermediate intensity levels correspond to two conformational states (S_H_ and S_I_) with two pores and one pore open, respectively. The low intensity level may represent a conformation (S_L_) with both pores closed.

**Fig. 2.**
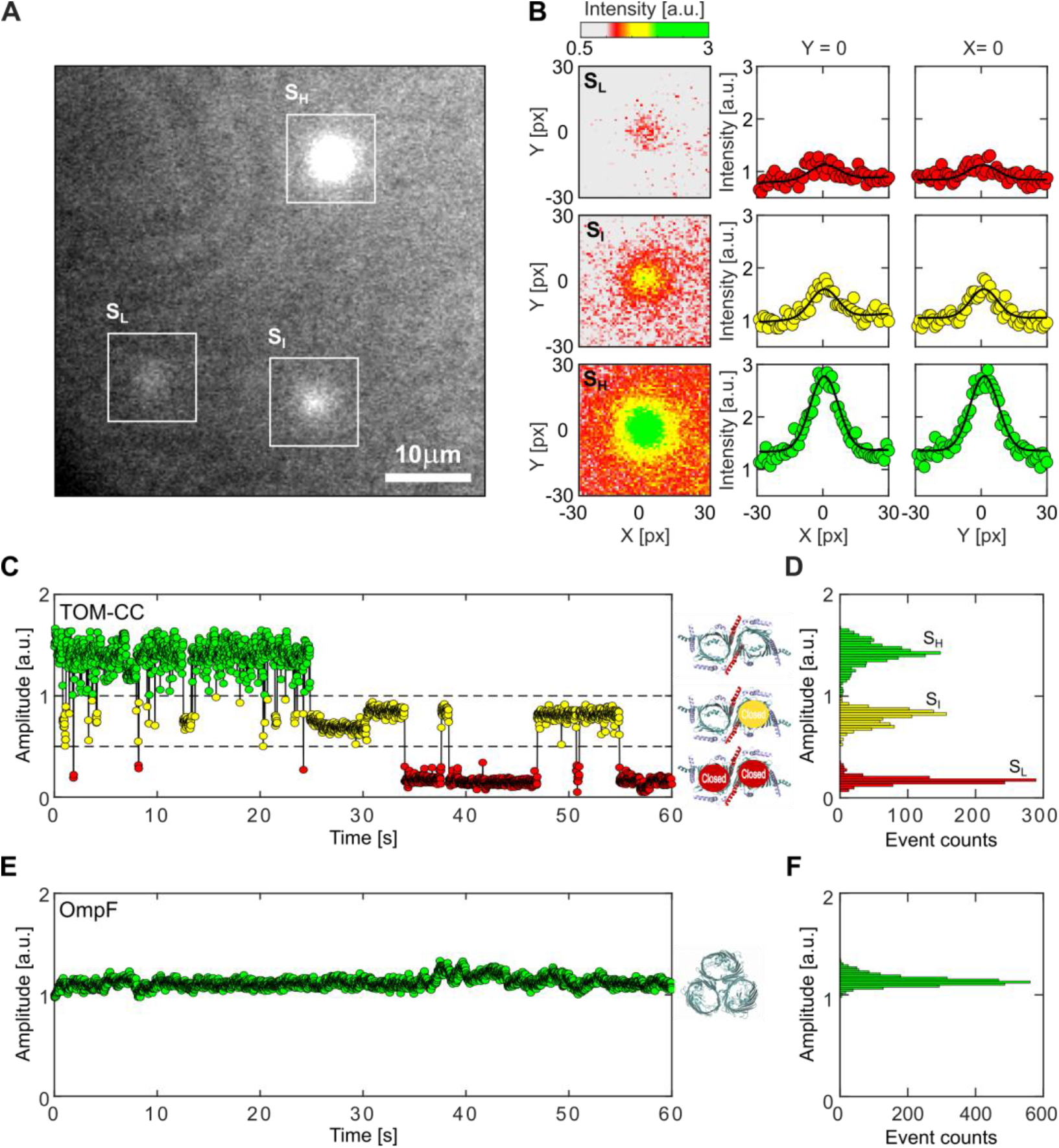
Visualizing the two pore channel activity of TOM-CC. (A) Typical image (*N* > 5.3 × 10^5^) of a non-modified agarose-supported DIB membrane with TOM-CC channels under 488 nm TIRF-illumination. The white squares mark spots of high (S_H_), intermediate (S_I_) and low (S_L_) intensity (Movie S1). (B) Fitting the fluorescence intensity profile of the three spots marked in (A) to two-dimensional Gaussian functions (Movie S2). Red, yellow and green intensity profiles represent TOM-CC in S_L_, S_I_ and S_H_ demonstrating Tom40 channels, which are fully closed, one and two channels open, respectively. Pixel size, 0.16 μm. (C) Fluorescence amplitude trace and (D) amplitude histogram of the two-pore β-barrel protein channel TOM-CC. The TOM-CC channel switches between S_H_, S_I_, and S_L_ permeability states over time. Inserts, schematic of *N. crassa* TOM core complex with two pores open in S_H_ (top), one pore open in S_I_ (middle), and two pores closed in S_L_ (bottom) (EMDB, EMD-3761; (Bausewein et al., 2020, 2017)); (E) Representative single-channel optical recording and (F) amplitude histogram of the three-pore β-barrel protein OmpF used as a control, which is completely embedded in the lipid bilayer. In contrast to the two-pore β-barrel protein complex TOM-CC, OmpF exhibits only one permeability state over time. Insert, model of *E. coli* OmpF (PDB, 1OPF) with all three pores open. Data were acquired by Ef-oSCR as described in Fig.1C at a frame rate of 47.5 s^−1^.

Movie S1 shows an optical recording of the open-closed channel activity of several TOM-CCs over time. Individual image frames of membranes were recorded at high frequency at a frame rate of 47.5 s^−1^ and corrected for fluorescence bleaching. The position and amplitude of individual spots were determined by fitting their intensity profiles to a two-dimensional symmetric Gaussian function with planar tilt to account for possible local illumination gradients in the bleaching corrected background (Fig. 2B and Movie S2). The time evolution of amplitude signals (Fig. 2C and Movie S2) shows that the TOM-CC does not occupy only one of the permeability states S_H_, S_I_ and S_L_, but can switch between these three permeability states over time (Fig. 2D). Additional examples are shown in Fig. S1. These data allow us to conclude that the fluctuations between the three defined permeability states are an inherent property of TOM-CC.

To rule out the possibility that the observed intensity fluctuations are caused by possible thermodynamical membrane undulations in the evanescent TIRF illumination field (Duncan et al., 2017), or by local variations in Ca^2+^ flux from *cis* to *trans* (Wang et al., 2018; Yuqin Wang et al., 2019), we visualized the ion flux through a three-pore β-barrel which is almost entirely embedded in the lipid bilayer (Figs. S2A and S2B). In a series of control experiments, we reconstituted *E. coli* OmpF (Benz, 2006) into DIB membranes and observed virtually constant fluorescence intensities (Figs. 2E and 2F, and Figs. S2C - S2F). The protein did not exhibit any gating transitions between its four different permeability states (one to three pores open, all pores closed) (Benz, 2006), which normally require voltage to induce closure. The toggling of the TOM-CC between the three different permeability states S_H_, S_I_ and S_L_ (Fig. 2C) therefore had to have another cause.

### The permeability states of TOM-CC are coupled to lateral mobility

Since the functions of many integral membrane proteins depend on their local position and state of movement in the membrane (Davis et al., 2021; Fujiwara et al., 2016; Jacobson et al., 2019; Koppel et al., 1981), we attempted to capture individual TOM-CCs in one of the permeability states S_H_, S_I_ or S_L_. To this end, we simultaneously tracked the open-closed activity and position of individual TOM-CC molecules in the membrane over time using the same experimental conditions as described above. A scheme of the experiment is shown in Fig. 3A.

**Fig. 3.**
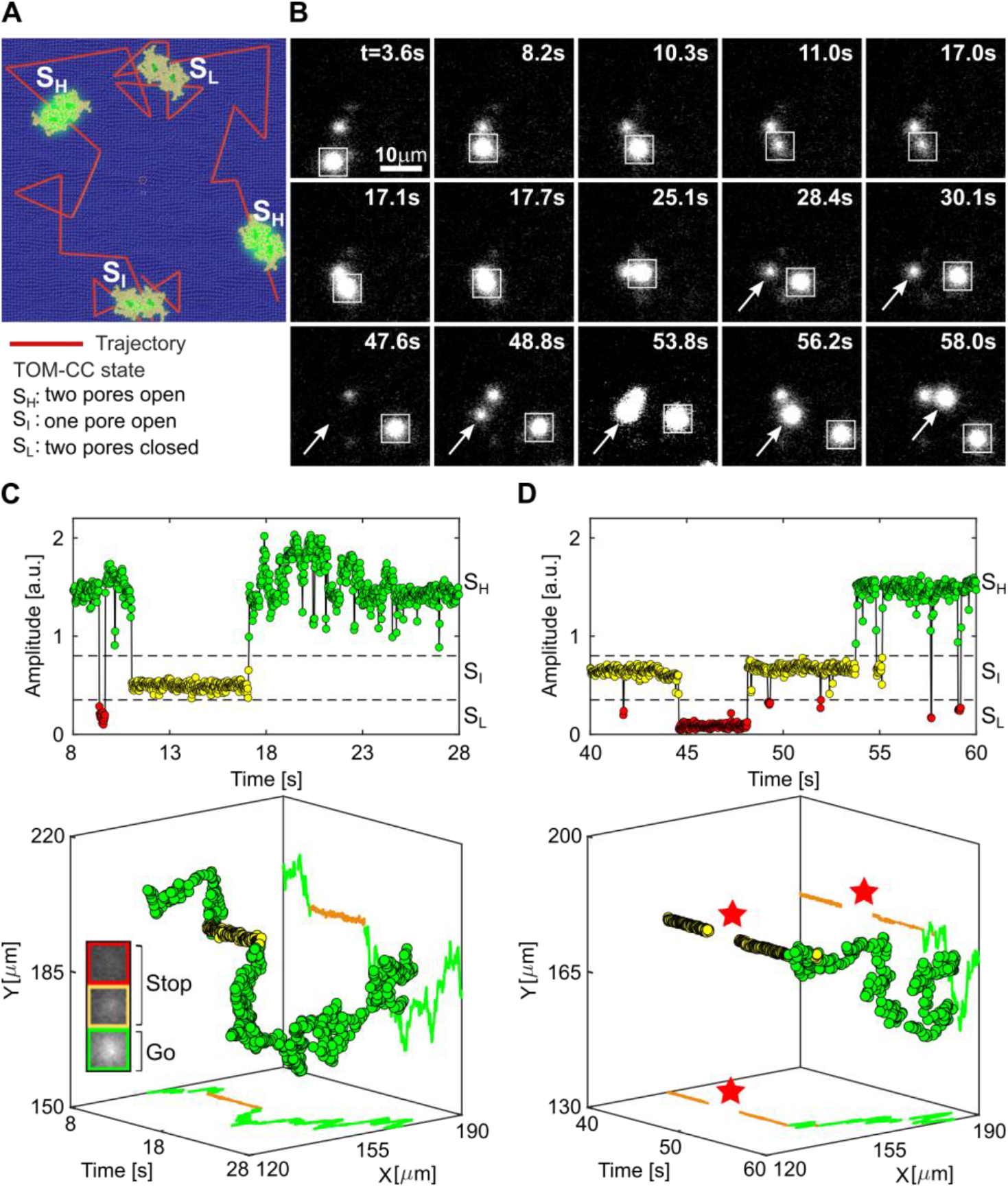
Lateral mobility correlates with the channel activity of TOM-CC. (A) Scheme for imaging both position and channel activity of single TOM-CCs. (B) Representative TIRF microscopy images of a non-modified agarose-supported DIB membrane with three TOM-CC molecules taken from a time series of 60 s. The square-marked spot displays lateral motion, interrupted by a transient arrest between *t* = 11.0 s and *t* = 17.0 s. The arrow-marked spot corresponds to a non-moving TOM-CC until *t* = 48.8 s. Afterwards, it starts moving. Both moving spots show high fluorescence intensity (S_H_); the non-moving spots display intermediate (S_I_) or dark (S_L_) fluorescence intensity (Movie S3). (C and D) Fluorescent amplitude trace and corresponding trajectory of the square- and arrow-marked TOM-CC as shown in (B) highlighted for two different time windows. Plots on top shows the change of amplitude over time, and plots on the bottom show the respective spatiotemporal dynamics for the three states. Comparison of the trajectories of single TOM-CC molecules with their corresponding amplitude traces reveals a direct correlation between *stop-and-go* movement and open-closed channel activity. Lateral diffusion of TOM-CCs in the DIB membrane is interrupted by temporary arrest, presumably due to transient linkage to the underlying agarose hydrogel. Although weak intensity profiles in S_I_ do not allow accurate position determination, the fluorescent spots disappear and reappear at the same spatial x,y coordinates (red stars). The higher amplitude (C, top) between *t* = 17.1 s and *t* = 25.1 s is due to the overlap between two adjacent spots. Green, TOM-CC is freely diffusive in S_H_; yellow and red, immobile TOM-CC in S_I_ and S_L_. Data were acquired by Ef-oSCR as described in Fig.1C at a frame rate of 47.5 s^−1^. A total of *n*_TOM_ = 64 amplitude traces and trajectories were analyzed.

Movie S3 and Fig. 3B clearly show that the open-closed channel activity of single TOM-CCs is coupled to lateral movement in the membrane. This is supported by comparison of the trajectories of single TOM-CC molecules with their corresponding fluorescence amplitude traces (Figs. 3C and 3D). The position of fluorescent spots does not change when TOM-CC is in intermediate S_I_ or low S_L_ permeability state. Although weak intensity profiles do not allow accurate determination of the position of TOM-CC in the membrane plane, Movie S3 and Fig. 3D clearly show that TOM-CC does not move in S_L_; disappearance and reappearance of the fluorescent spot, switching from S_I_ to S_L_ and back to S_I_, occurs at virtually the same spatial x,y-coordinates. In contrast, the trajectories of TOM-CC in S_H_ state demonstrate free diffusion. Additional samples of trajectories and amplitude traces are shown in Fig. S1.

Similar *stop-and-go* movement patterns were observed in an independent set of experiments for TOM-CC covalently labeled with fluorescent dye Cy3 (Movie S4 and Fig. S3). It is particularly striking that freely moving TOM-CC molecules stop at the same spatial x,y-position when they cross the same position a second time, indicating a specific molecular trap or anchor point at this stop-position below the membrane. OmpF, however, used as a control, shows the most elementary mode of mobility expected for homogeneous membranes: simple Brownian translational diffusion (Movie S5 and Fig. S2). Since OmpF is almost entirely embedded in the membrane, there seems to be no coupling between channel activity and lateral protein diffusion.

In good agreement with these results, the diffusion coefficients of the TOM-CC, evaluated from the activity profiles and trajectories (Fig. 3 and Fig. S1), were determined from time-averaged mean squared displacements as *D*_I_ = *D*_L_ ≤ *D*_min_ = 0.01 μm^2^s^−1^ and *D*_H_ ≃ 0.85 ± 0.16 (*mean* ± *SEM, n* = 46) in states S_I_ and S_H_, respectively. TOM-CC molecules, that revealed a diffusion constant less or equal than *D*_min_, were defined as immobilized. The diffusion coefficient *D*_H_ corresponds to typical values of Tom40 (*D*_Tom40_ ∼0.5 μm^2^ s^−1^) and Tom7 (*D*_Tom7_ ∼0.7 μm^2^ s^−1^) in native mitochondrial membranes (Kuzmenko et al., 2011; Sukhorukov et al., 2010) and is comparable to that of transmembrane proteins in plasma membranes lined by cytoskeletal networks (*39, 43*). The diffusion coefficient of TOM-CC in S_L_ state could not always be reliably determined here due to its extremely low intensity levels (see Materials and Methods).

TOM-CC labeled with Cy3 yielded diffusion constants of *D*_Cy3,H_ ≃ 0.36 ± 0.08 μm^2^ s^−1^ (mean ± SEM, *n* = 15) and *D*_Cy3,I_ ≤ *D*_min_ = 0.01 μm^2^s^−1^ for moving and transiently trapped particles, respectively (Fig. S3). In line with typical values for protein in homogenous lipid membranes (Koppel et al., 1981; Ramadurai et al., 2009), the lateral diffusion coefficient of the control protein OmpF was determined as *D*_OmpF_ ≃ 1.16 ± 0.07 μm^2^ s^−1^ (mean ± SEM, *n* = 42) (Fig. S2).

Based on these results we conclude that transient arrest of TOM-CC in a lipid bilayer membrane, caused by interaction with the supporting hydrogel, triggers the closure of its two β-barrel pores. Since retarded diffusion of membrane proteins is largely the result of cluster formation of proteins near membrane surfaces (Nawrocki et al., 2019), this phenomenon is important for studying preprotein trapping at either Tom40 β-barrel when TOM-CC interacts with proteins at the periphery of the outer membrane.

### Controlled immobilization of the TOM-CC results in channel closures

Because the two Tom22 subunits in the middle of the TOM-CC clearly protrude from the membrane plane at their IMS side (Fig. 1B) (Araiso et al., 2019; Bausewein et al., 2017; Tucker and Park, 2019; Wang et al., 2020), we hypothesized that Tom22 functions as a “light-switch” and determines the lateral mobility of the TOM-CC and thus the transitions between open (S_H_) and the closed (S_I_ and S_L_) conformations of the two TOM-CC pores.

To test this hypothesis, we replaced the non-modified agarose hydrogel with Ni-NTA-modified agarose to further restrict lateral movement of the protein by permanently tethering individual TOM-CC molecules via the C-terminus of histidine-labeled Tom22 to the hydrogel (Fig. 4A).

**Fig. 4.**
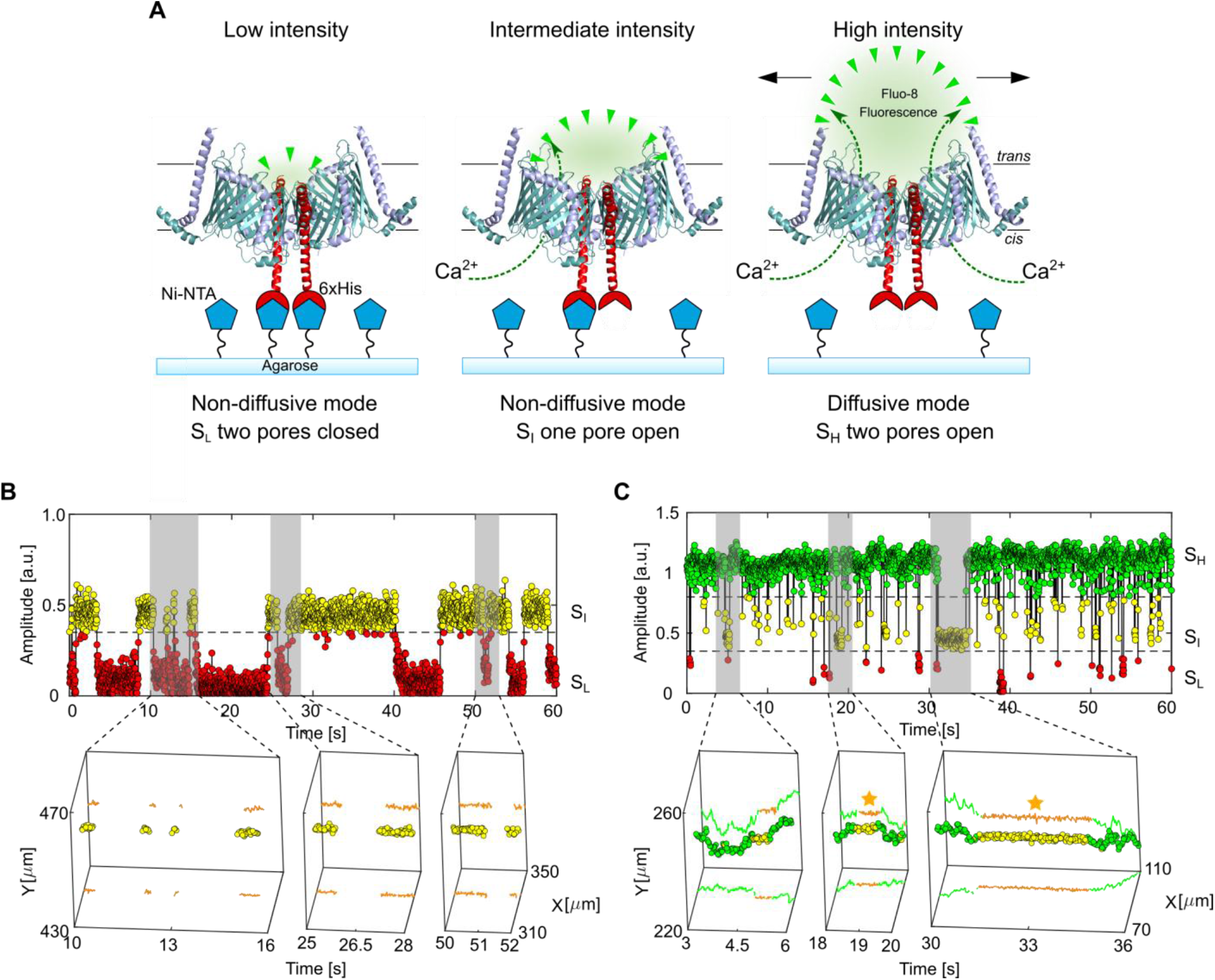
Controlled immobilization of TOM-CC triggers channel closures. (A) Schematic representation of individual TOM-CC channels in DIB membranes supported by Ni-NTA-modified agarose. TOM-CC molecules can be permanently linked to the underlying hydrogel via His-tagged Tom22. Tethered and non-tethered TOM-CC molecules in closed (S_I_ and S_L_) and open (S_H_) states are indicated, respectively. (B) Fluorescent amplitude trace (top) of a TOM-CC channel permanently tethered to Ni-NTA-modified agarose. The trajectory segments (bottom) correspond to the time periods of the amplitude traces marked in grey (Movie S6: bottom). Non-diffusive, permanently immobilized TOM-CC is only found in S_I_ or S_L_, indicating that tight binding of the His-tagged Tom22 domain (Fig. 1B) to Ni-NTA-modified agarose triggers closure of the β-barrel TOM-CC pores. (C) Fluorescence amplitude trace (top) of a TOM-CC channel transiently and non-specifically entangled by Ni-NTA-modified agarose. The trajectory segments (bottom) correspond to the time periods of the amplitude traces marked in grey (Movie S6 bottom). The movement of TOM-CC is interrupted twice at the same spatial *x,y* membrane position from *t*_1_ = 18.56 s to *t*_2_ = 19.19 s and from *t*_3_ = 31.14 s to *t*_4_ = 34.55 s (yellow stars). Consistent with the data shown in Fig. 3, moving TOM-CC molecules in diffusive mode are found in the fully open S_H_ state; transient tethering causes the TOM-CC β-barrels to close. Data were acquired by Ef-oSCR as described in Fig.1C at a frame rate of 47.5 s^−1^. A total of *n*_TOM_ = 123 amplitude traces and trajectories were analyzed.

In good agreement with the observation that transient closure of the two Tom40 β-barrels accompanies transient arrest of the TOM-CC (Figs. 3C and 3D), permanently immobilized TOM-CC (*D*_I, L_ ≤ *D*_min_ = 0.01 μm^2^ s^−1^, *n* = 83) was most often found in states S_I_ or S_L_, indicating one or two pores closed, respectively (Movie S6 and Fig. 4B, Fig. S4A - S4D). TOM-CC molecules that were not immobilized by the binding of Tom22 to Ni-NTA moved randomly in the membrane plane (*D*_H_ ≃ 0.34 ± 0.06 μm^2^ s^−1^, *mean* ± *SEM, n* = 40). As expected, movement was interrupted occasionally by periods of transient arrest (*D*_I, L_ ≤ *D*_min_). Again, moving TOM-CC molecules were found in the fully open S_H_ state; non-moving complexes in the S_I_ or the S_L_ states (Fig. 4C; Figs. S4E - S4H). In the presence of imidazole, which prevents tight binding of histidine-tagged Tom22 to Ni-NTA-modified agarose, virtually no permanently immobilized TOM-CC molecules were observed (*D*_H_ ≃ 1.35 ± 0.14 μm^2^ s^−1^, *mean* ± *SEM, n* = 15).

### Correlation between lateral motion and TOM-CC channel activity

DIB membranes supported by both hydrogels, non-modified and Ni-NTA-modified agarose (Movie S6), showed a statistically significant number of single TOM-CC molecules that were either non-diffusive *D*_H, I_ ≤ *D*_min_ or diffusive *D*_H_ > *D*_min_ at S_I_ and S_H_. Thus, TOM-CC molecules were numerically sorted into diffusive and permanently tethered groups by *D*_H_ and *D*_I_ to emphasis the correlation between the mode of lateral diffusion and channel activity of 187 observed TOM-CC molecules (Fig. 5A). We can generally define three different classes of lateral motion and channel activity. The first and major class (I) shows lateral mobility at S_H_ only, while being tethered at S_I_ (*D*_H_ > *D*_min_ and *D*_I_ ≤ *D*_min_). The second class (II) shows similar diffusivities (*D*_H, I_ > *D*_min_, Fig. S5) at both states, S_H_ and S_I_. The TOM molecules of this class probably do not have functional Tom22 (Ahting et al., 2001; Shiota et al., 2015) and thus do not interact with the hydrogel. Another possible but unlikely explanation is a spatial void of agarose network preventing mechanoregulated interaction of Tom22 with the network within the observation time window. The third class (III) represents permanently tethered TOM-CC molecules, which are exclusively non-diffusive (*D*_H, I_ ≤ *D*_min_). Most molecules of that class are in S_I_ and S_L_ (Figs. S4A and S4B). Those TOM-CC molecules, which briefly change from S_I_ to S_H_ and back to S_I_ (Figs. S4C and S4D), might become diffusive but are immediately recaptured and trapped again by the hydrogel below the membrane.

**Fig. 5.**
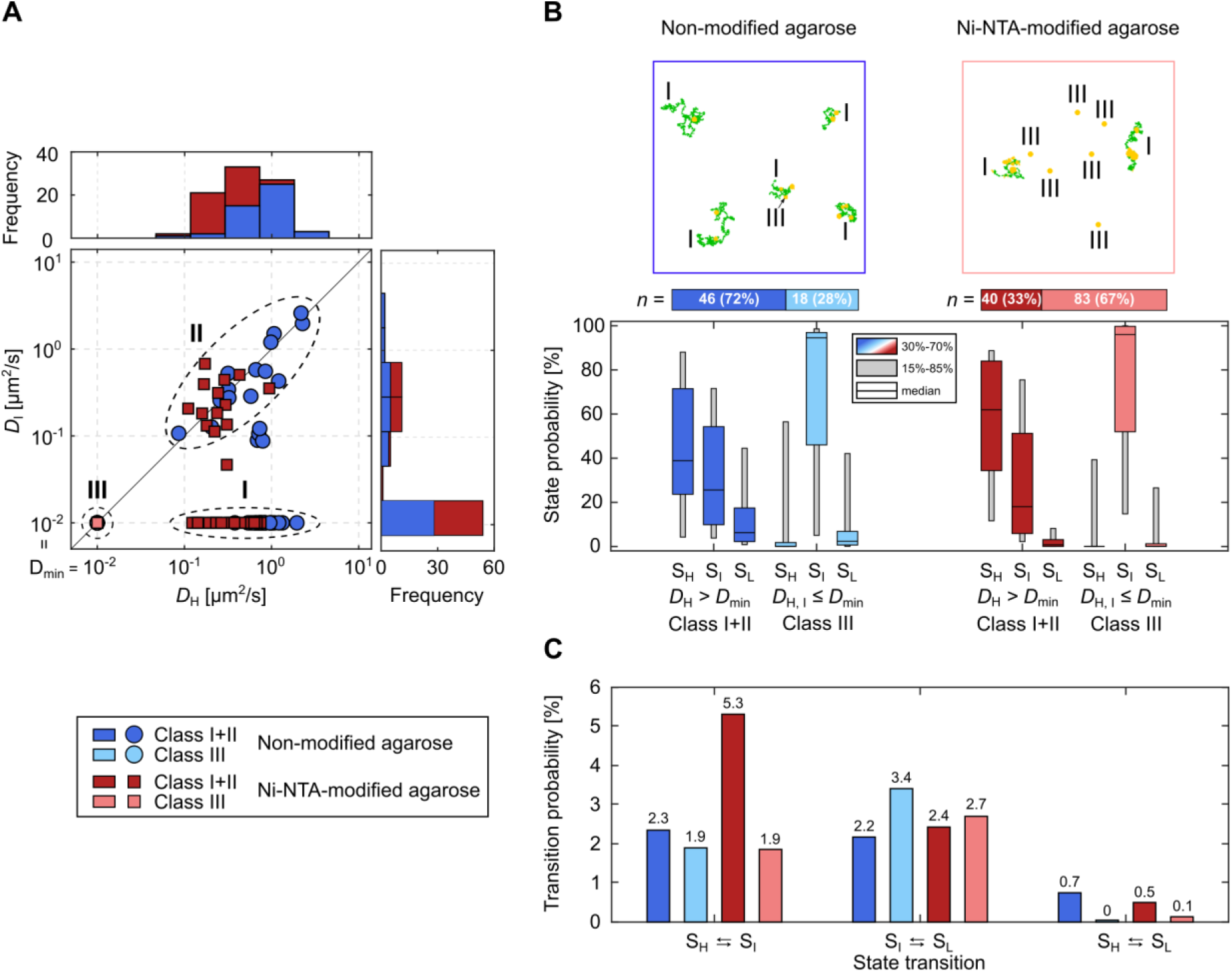
Statistical correlation between channel activity and lateral mobility of TOM-CC. (A) *D*_I_ as a function of *D*_H_ individually plotted for all TOM-CC molecules in DIB membranes supported by non-modified (dark blue and light blue, *n* = 64) and Ni-NTA modified agarose (dark red and light red, *n* = 123). Frequency histograms of *D*_H_ and *D*_I_ are shown on top and right side, respectively. Three classes can be defined: a main class of moving particles in S_H_ while being transiently tethered at S_I_ (I), a second class of freely moving particles in S_H_ and S_I_ (II) and a third class of permanently tethered molecules in S_I_ and S_L_ (III). (B) Example trajectories (top and Movie S6) and state probabilities (bottom) of non-permanently (*D*_H_ > *D*_min_, class I+II) and permanently tethered (*D*_H, I_ ≤ *D*_min_, class III) TOM-CC in DIBs supported by non-modified and Ni-NTA-modified agarose. The probability of being in state S_H_ is higher for non-permanently (*D*_H_ > *D*_min_, class I+II, dark blue [*n* = 46] and dark red [*n* = 40]) than for permanently tethered molecules (*D*_H, I_ ≤ *D*_min_, class III, light blue [*n* = 18] and light red [*n* = 83]). The probability of being in state S_I_ is significantly higher for permanently (*D*_H, I_ ≤ *D*_min_, class III) than for non-permanently tethered particles (*D*_H_ > *D*_min_, class I+II). This suggests that binding of Tom22 to Ni-NTA agarose below the membrane triggers closure of the TOM-CC channels. The data are represented as median; the confidence intervals are given between 15% to 85% and 30% to 70%. Moving particles in S_H_ are shown in the trajectories in green; transiently or permanently tethered molecules in S_I_ are shown in yellow. (C) Absolute state transition probabilities classified by bidirectional state transitions as S_H_ ⇆ S_I_, S_I_ ⇆ S_L_, and S_H_ ⇆ S_L_. Diffusive TOM-CC molecules have a significantly higher transition probability for switching between S_H_ and S_I_ in DIBs supported by Ni-NTA-modified agarose (∼5.3%) than in non-modified agarose membranes (∼2.3%). This is consistent with the higher efficacy of TOM-CC-trapping by Ni-NTA-modified agarose compared to non-modified agarose. Classification of non-permanently and permanently tethered TOM-CC is shown at the left bottom.

Fig. 5B shows state probabilities of TOM-CC in membranes supported by the two different hydrogels, non-modified and Ni-NTA-modified agarose. Diffusive molecules (*D*_H_ > *D*_min_) in membranes supported both by non-modified and Ni-NTA-modified agarose show similar probabilities to be at one of the three permeability states (S_H_, S_I_ and S_L_). Diffusive TOM-CC molecules are significantly more often at S_H_ than at S_I_ and S_L_. The permanently tethered fraction of TOM-CC (67%) in Ni-NTA-modified agarose is ∼2.4 times larger compared to the fraction (28%) in non-modified agarose, consistent with the stronger interaction of Tom22 with the hydrogel, thereby permanently constraining lateral motion. In line with this, permanently tethered molecules (*D*_H, I_ ≤ *D*_min_) in both hydrogel-supported membranes stay at S_I_ during the majority of time, and show only transient S_H_ and S_L_ occupancy. The data suggest that the C-terminal IMS domain of Tom22 (Fig. 1B) plays a previously unrecognized role in mechanoregulation of TOM-CC channel activity by binding to immobile structures near the membrane.

Although diffusive TOM-CC molecules (*D*_H_ > *D*_min_) are observed more often at S_H_ in Ni-NTA-modified agarose than in non-modified agarose supported membranes (Fig. 5B), they show a lower stability at S_H_ having a significantly higher transition probability for switching between S_H_ and S_I_ (Fig. 5C, (S_H_ ⇆ S_I_) ≃ 5.3% versus (S_H_ ⇆ S_I_) ≃ 2.3%). This is in line with the higher efficacy of TOM-CC-trapping by Ni-NTA-modified agarose compared to non-modified agarose. In contrast to non-modified hydrogel, Ni-NTA-modified agarose hydrogel can capture freely mobile TOM-CC via the IMS domain of Tom22 in two ways: by specific interaction and permanent anchorage with Ni-NTA and by nonspecific collision and transient anchorage. While a direct transition between S_H_ and S_L_ barely occurs in both systems, transitions between S_I_ and S_L_ are similarly often. This indicates that the two Tom40 β-barrel pores independently open and close within the time resolution (∼20 ms) of our experiment.

## Discussion

TOM-CC is usually considered as a passive conduit for mitochondrial preproteins and small molecules across the outer mitochondrial membrane, which is regulated by interactions with other proteins and posttranslational modifications (Rapaport et al., 1998; Schmidt et al., 2011).

In this study, we have shown for the first time that lateral diffusion of isolated TOM-CC in the plane of the membrane is coupled to transmembrane channel activity. Conversely, restriction of the lateral mobility of the complex by the surrounding matrix influences channel activity in a defined manner. This in turn indicates that TOM-CC does not only respond to biochemical (Rapaport et al., 1998; Schmidt et al., 2011), but also to yet-unknown mechanical stimuli: anchorage of freely moving TOM-CC to structures near the membrane leads to TOM-CC channel closing.

Our conclusions are based upon data obtained from single molecules, which yield insights into molecular events that are difficult to obtain *in vivo* and in bulk methods. Typically, single molecule methods are often subject to statistical uncertainties arising from the limited amount of measurements, which are technically feasible. For this reason, we have obtained data from *n* = 187 single molecules and over half a million image frames (Fig. 5A, *N* = 532,576). This sampling number allows us to assign molecular events with statistical confidence using non-parametric statistics, but even a visual perusal of the data is sufficient to allow a first assessment of data reliability. Our data indicate three molecular states of the TOM-CC, for which the simplest model is a dimeric pore structure with one or two pores open and all pores closed. As indicated above, the state transitions are responding to interaction with the supporting agarose matrix. We believe this mechanoregulated interactive effect, as we observed, is of physiological relevance.

Previous super-resolution fluorescence and immuno-electron microscopy studies have revealed a dynamic submitochondrial distribution of the protein translocation machineries TOM and TIM23 in the outer and inner membrane of mitochondria, both of which depend on the physiological state of the cell (Palmer et al., 2021; Vogel et al., 2006; Wurm et al., 2011). An attractive possibility would be that interaction of the TOM complex with subunits of TIM23 in the mitochondrial intermembrane space (IMS) *via* their subunits Tom22 and Tim50, respectively, might be coordinated so that protein import through the TOM channels can only occur when TIM23 is locally available (Callegari et al., 2020; Chacinska et al., 2005; Mokranjac et al., 2009; van der Laan et al., 2006). Thus, in the light of transient cooperation between TOM and TIM23 (Donzeau et al., 2000; Gevorkyan-Airapetov et al., 2009; Waegemann et al., 2015) it is becoming increasingly clear that the interplay between lateral diffusion and channel activity of these complexes is pivotal for protein translocation into mitochondria.

However, previous studies beg a fundamental question: are the two Tom40 channels open continually, as seems to be the case from the static cryoEM structures (Fig. 1B; (Araiso et al., 2019; Bausewein et al., 2017; Tucker and Park, 2019; Wang et al., 2020), or can the protein channels be actively influenced by interaction with exogenous, interacting proteins? Previous studies reported that TOM complex channels from wild-type yeast mitochondria were mainly in the closed state (van Wilpe et al., 1999). However, TOM isolated from mitochondria lacking Tom22 was mainly in the open state (van Wilpe et al., 1999).

In our study, we have demonstrated that Tom22 plays an important role in the kinetics of TOM-CC channel activation. Examination of the cryoEM structure of *N. crassa* TOM-CC (Fig. 1B, (Bausewein et al., 2020, 2017)) shows that its two Tom22 subunits extend significantly into the IMS space, thereby preventing a direct interaction of the Tom40 β-barrels with the agarose matrix. It is therefore likely that only the two Tom22 subunits are primarily responsible for the matrix-dependent mechanoregulated channel closing activity. First, this hypothesis is supported by our observation that His-tagged Tom22, when tightly bound by Ni-NTA-agarose, triggers channel closing (Movie S6). Secondly, the cryoEM structure of TOM-CC reveals that the dimer interface of the Tom40 barrels is extremely limited, with only nine amino acid residues contributing to the interface at the cytosolic side of the barrel (Araiso et al., 2019; Bausewein et al., 2020, 2017; Tucker and Park, 2019; Wang et al., 2020). At the IMS side of the barrel, two conserved lysine residues (Tom40-K298, contributed from each *Nc. crassa* Tom40 monomer) seem to be close enough to severely destabilize the dimer. In fact, most of the stabilizing interactions of the Tom40 dimer seem to be contributed by residues from the interacting, transmembrane α-helices of Tom22 and a lipid phosphate group (Araiso et al., 2019; Bausewein et al., 2020; Tucker and Park, 2019). Thus, it seems reasonable that the conformation of the Tom40 barrel dimer is essentially determined by the positioning of two Tom22 subunits. Taken together, it is tempting to suggest that, in our experiments, when the intermembrane space (IMS)-located polar domains of the Tom22 subunits (which extend more than 22Å into the IMS, Fig. 1B) interact with the matrix of the agarose gel, the Tom22 subunits force the Tom40 dimer interface to experience a conformational change resulting in channel closure.

From a mechanistic point of view, this might be caused by classic rigid body rotation at the Tom40 dimeric interface, perpendicular to the membrane normal. Rigid body rotation at subunit interfaces is a common feature of quaternary conformational changes in oligomeric proteins, for which the archetypical paradigm is rotation of the α_1_β_1_−α_2_β_2_ interface of hemoglobin during the oxy-deoxy allosteric transition (Baldwin and Chothia, 1979). The Tom40 dimer rigid body rotation might allow the disordered 84 amino acid residues, cytosolically located N-terminal domain of the Tom22 subunit, which is not visible in the cryoEM structures, to “fall into” the Tom40 channel, thereby blocking Ca^2+^ export in the TIRF experiment. *In vivo*, this conformational change might be caused by tethering the IMS domain of Tom22 to Tim50 of the TIM23 machinery, which is virtually immobile in the mitochondrial inner boundary membrane (*D*_Tim23_ ≃ 0.04 μm^2^s^−1^) (Albrecht et al., 2006; Appelhans and Busch, 2017; Callegari et al., 2020; Chacinska et al., 2005; Mokranjac et al., 2009; van der Laan et al., 2006).

Previous studies suggested that a dimeric Tom22-free form of Tom40 dynamically exchanges with a trimeric TOM complex in outer mitochondrial membranes (Araiso et al., 2019; Shiota et al., 2015). In this study, we have not explicitly considered the functional relevance of TOM trimers. First, all of the cryoEM data obtained for detergent-solubilized TOM-CCs published so far (Araiso et al., 2019; Bausewein et al., 2017; Tucker and Park, 2019; Wang et al., 2020), indicates the dimer as the major subunit form. Secondly, we do not require a trimer to interpret the channel activity observed here. In fact, the functional trimer would require a fourth channel state, which has rarely been observed. Finally, the formation of trimers from functional dimers seems to be at variance with well-known principles of protein symmetry at subunit interfaces. Therefore, in lieu of more biophysical data from defined systems, we presently do consider the dimer and not the trimer of Tom40 to be the major functional form of the TOM-CC.

While eukaryotic secretion systems allow passage through a single bilayer, proteins that enter organelles of endosymbiotic origin must pass through two bilayers and translocation machineries to reach the interior of the organelle (Davis et al., 2021). Thus, future approaches should consider the multiprotein nature of the TOM-TIM import system, and how this additional layer of complexity is affected by the role of mechanoregulation in native mitochondrial membranes with TOM subunits Tom20 and Tom70 in the presence of mitochondrial preproteins (Wiedemann and Pfanner, 2017). However, since Tom20 and Tom70 do not have appreciable IMS domains and both proteins are only loosely bound to the complex (Ahting et al., 1999; Wiedemann and Pfanner, 2017), it seems unlikely that these proteins significantly influence the interplay between lateral diffusion and channel kinetics of the TOM-CC. The diffusion coefficient of mobile TOM-CC shown in our work is in good agreement to that of mobile Tom40 in native mitochondria (Kuzmenko et al., 2011; Sukhorukov et al., 2010). We therefore propose that the *stop-and-go* concept described here also applies to native mitochondria, in which proteins can freely move only on smaller length scales. TOM-CC channels may simply be trapped more frequently, leading to a higher frequency of open-closed transitions.

Recently, the now extensive structural and physical data common to mechanosensitive membrane proteins have been reviewed (Haswell et al., 2011; Jin et al., 2020; Kefauver et al., 2020). The structural data now allows mechanosensitive proteins (all of which were comprised of transmembrane α-helices) to be separated into five different classes, each subject to characteristic underlying molecular mechanisms. They also show that for the membrane channels considered, fundamental physical properties of the membrane can influence channel activity. This ancient mechanism to regulate open-closed channel activity is also observed in mitochondria (Deng et al., 2020; Lee et al., 2016; Li et al., 2020; Walewska et al., 2018).

Mechanosensitivity *via* force transmission from a tether to the extracellular matrix and/or the cytoskeleton has been established for a number of α-helical membrane proteins (Brohawn et al., 2014a, 2014b; Ge et al., 2018; Jin et al., 2020; Kefauver et al., 2020; Martinac, 2004; Pliotas and Naismith, 2017; Teng et al., 2015; Yang Wang et al., 2019). These findings go along with our observations for the TOM-CC, which seems to obey the “tether model” proposed by Kefauver et al. (2020) (Kefauver et al., 2020), where an intermembrane protein anchor domain limits lateral diffusion. To our knowledge, the TOM-CC complex would thus be the only example so far, of a membrane β-barrel protein, which exhibits membrane state-dependent mechanosensitive-like properties.

## Materials and Methods

### Growth of *Neurospora crassa* and preparation of mitochondria

*Neurospora crassa* (strain GR-107) that contains a hexahistidinyl-tagged form of Tom22 was grown and mitochondria were isolated as described (Künkele et al., 1998). Briefly, ∼1.5 kg (wet weight) of hyphae were homogenized in 250 mM sucrose, 2 mM EDTA, 20 mM Tris pH 8.5, 1 mM phenylmethylsulfonyl fluoride (PMSF) in a Waring blender at 4°C. ∼1.5 kg of quartz sand was added and the cell walls were disrupted by passing the suspension through a corundum stone mill. Cellular residues were pelleted and discarded in two centrifugation steps (4,000 x g) for 5 min at 4°C. The mitochondria were sedimented in 250 mM sucrose, 2 mM EDTA, 20 mM Tris pH 8.5, 1 mM PMSF at 17,000 x g for 80 min. This step was repeated to improve the purity. The isolated mitochondria were suspended in 250 mM sucrose, 20 mM Tris pH 8.5, 1 mM PMSF at a final protein concentration of 50 mg ml^−1^, shock-frozen in liquid nitrogen and stored at −20°C.

### Isolation of TOM core complex

TOM-CC, containing subunits Tom40, Tom22, Tom7, Tom6 and Tom5, were purified from *Neurospora crassa* strain GR-107 as described (Bausewein et al., 2017). *N. crassa* mitochondria were solubilized at a protein concentration of 10 mg/ml in 1% (w/v) n-dodecyl-β-D-maltoside (Glycon Biochemicals, Germany), 20% (v/v) glycerol, 300 mM NaCl, 20 mM imidazole, 20 mM Tris-HCl (pH 8.5), and 1 mM PMSF. After centrifugation at 130,000 x g, the clarified extract was loaded onto a nickel-nitrilotriacetic acid column (Cytiva, Germany). The column was rinsed with the same buffer containing 0.1% (w/v) n-dodecyl-β-D-maltoside and TOM core complex was eluted with buffer containing 0.1% (w/v) n-dodecyl-β-D-maltoside, 10 % (w/v) glycerol, 20 mM Tris (pH 8.5), 1 mM PMSF, and 300 mM imidazole. For further purification, TOM core complex containing fractions were pooled and loaded onto a Resource Q anion exchange column (Cytiva) equilibrated with 20 mM Hepes (pH 7.2), 2% (v/v) dimethyl sulfoxide and 0.1 % (w/v) n-dodecyl-β-D-maltoside. TOM core complex was eluted with 0 – 500 mM KCl. A few preparations contained additional phosphate (∼0.19 mM). The purity of protein samples (0.4 – 1.2 mg/ml) was assessed by sodium dodecyl sulfate polyacrylamide gel electrophoresis (SDS-PAGE) followed by staining with Coomassie Brilliant Blue.

### Fluorescence labeling of TOM core complex

TOM-CC was covalently labeled with the fluorescent dye Cy3 according to (Joo and Ha, 2008). Briefly, about 1 mg/ml of purified TOM-CC was reacted with Cy3-maleimide (AAT Bioquest, USA) at a molar ratio complex to dye of 1:5 in 20 mM HEPES (pH 7.2), 2 % (v/v) dimethyl sulfoxide, 350 mM KCl and 0.1 % (w/v) n-dodecyl-β-D-maltoside at 25 °C for 2 h in the dark. Labeled protein was separated from unconjugated dye by affinity chromatography using Ni-NTA resin, subjected to SDS-PAGE and visualized by 555 nm light and Coomassie Brilliant Blue staining.

### Isolation of OmpF

Native OmpF protein was purified from *Escherichia coli* strain BE BL21(DE3)omp6, lacking both LamB and OmpC as described (Bieligmeyer et al., 2016). Cells from 1 L culture were suspended in 50 mM Tris-HCl, pH 7.5 buffer containing 2 mM MgCl_2_ and DNAse and broken by passing through a French press. Unbroken cells were removed by a low-speed centrifugation, then, the supernatant was centrifuged at 100,000 x g for 1 h. The pellet was resuspended in 50 mM Tris-HCl, pH 7.5, and mixed with an equal volume of SDS buffer containing 4 % (w/v) sodium dodecyl sulfate (SDS), 2 mM β-mercaptoethanol and 50 mM Tris-HCl, pH 7.5. After 30 min incubation at a temperature of 50 °C, the solution was centrifuged at 100,000 x g for 1 h. The pellet was suspended in 2 % (w/v) SDS, 0.5 M NaCl, 50 mM Tris-HCl, pH 7.5, incubated at a temperature of 37 °C for 30 min and centrifuged again at 100,000 x g for 30 min. The supernatant containing OmpF was dialyzed overnight against 20 mM Tris, pH 8, 1 mM EDTA and 1 % (w/v) n-octyl polyoxyethylene (Octyl-POE, Bachem, Switzerland). The purity of the protein was assessed by SDS-PAGE.

### Formation of droplet interface bilayers

Droplet interface bilayer (DIB) membranes were prepared as previously described (Huang et al., 2015; Leptihn et al., 2013; Wang et al., 2018; Yuqin Wang et al., 2019) with minor modifications. Glass coverslips were washed in an ultrasonic bath with acetone. Then the coverslips were rinsed several times with deionized water and dried under a stream of nitrogen. Subsequently, the glass coverslips were subjected to plasma cleaning for 5 min. 140 μl of molten 0.75% (w/v) low melting non-modified agarose (T_m_ < 65°C, Sigma-Aldrich) or alternatively low melting Ni-NTA-modified agarose (Cube Biotech, Germany) was spin-coated at 5,000 rpm for 30 s onto the plasma-cleaned side of a glass coverslip. After assembly of the coverslip in a custom-built DIB device, the hydrogel film was hydrated with 2.5 % (w/v) low melting agarose, 0.66 M CaCl_2_ and 8.8 mM HEPES (pH 7.2) or with 2.5 % (w/v) low melting agarose, 0.66 M CaCl_2_, 300 mM imidazole and 8.8 mM HEPES (pH 7.2) and covered with a lipid/oil solution containing 9.5 mg/ml 1,2-diphytanoyl-sn-glycero-3-phosphocholine (DPhPC, Avanti Polar Lipids, USA) and 1:1 (v:v) mixture oil of hexadecane (Sigma-Aldrich) and silicon oil (Sigma-Aldrich). Aqueous droplets (∼200 nl) containing 7 μM Fluo-8 sodium salt with a maximum excitation wavelength of 495 nm (Santa Cruz Biotechnology, USA), 400 μM EDTA, 8.8 mM HEPES (pH 7.2), 1.32 M KCl, ∼2.7 nM TOM core complex or ∼2 nM OmpF were pipetted into the same lipid/oil solution in a separate tray using a Nanoliter 2000 injector (World Precision Instruments, Sarasota, USA). After 2 hours of equilibration at room temperature, the droplets were transferred into the custom-built DIB device to form stable lipid bilayers between the droplet and the agarose hydrogel.

### TIRF microscopy and optical recording

An inverted total internal reflection fluorescence microscope (Ti-E Nikon) was used to image DIB membranes under TIRF illumination using a 488 nm laser (Visitron). Fluorescent emission of Fluo-8, transmitted through a Quad-Band TIRF-Filter (446/523/600/677 HC, AHF), was collected through a 100 x oil objective lens (Apochromat N.A. 1.49, Nikon) and recorded by a back-illuminated electron-multiplying CCD camera (iXon Ultra 897, 512 × 512 pixels, Andor) for 1 min at a frame rate of 47.51 s^−1^. The pixel size was 0.16 μm.

### Tracking of fluorescence spots

For reliable tracking and analysis of the spatiotemporal dynamics of individual fluorescence TOM-CC channel activity a customized fully automated analysis routine was implemented in Matlab (The Mathworks, USA). The effect of bleaching was corrected by considering a double exponential decay obtained by least square fitting of the image series. No filter algorithm was applied. The initial spatial position of a fluorescence spot was manually selected and within a defined region of interested (ROI, 30 × 30 pixels) fitted to a two-dimensional symmetric Gaussian function with planar tilt that accounts for possible local illumination gradients, as follows

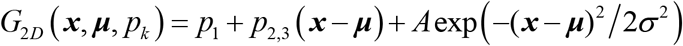

where ***x*** *=* (*x, y*) is the ROI with the fluorescence intensity information, A and σ are the amplitude and width of the Gaussian, *p*_*k*_ are parameters that characterize the background intensity of the ROI, and ***μ*** *=* (*x*_0_, *y*_0_) defines the position of the Gaussian. The latter was used to update the position of ROI for the next image. Spots that temporal fuse their fluorescence signal with closely located spots were not considered due to the risk of confusing those spots.

### Data analysis

The extracted amplitudes were separated by two individually selected amplitude-thresholds that dived the three states of activity (S_H_, S_I_ and S_L_). The lateral diffusion constants *D*_H_ and *D*_I_ were obtained individually for spots within the respective high and intermediate amplitude range by linear regression of the time delay *τ* and the mean square displacement of the spots. Further details may be found in SI Experimental Methods. As

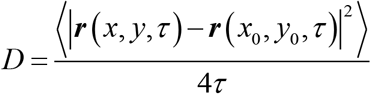

The largest time delay *τ*_max_ = 0.5 s was iteratively decreased to suffice the coefficient of determination *R*^2^ ≥ 0.9 for avoiding the influence of sub-diffusion or insufficient amount of data. This followed the general approach of a Brownian particle in two-dimensions. The diffusions less than *D*_min_ = 0.01 μm^2^ s^−1^ are defined as non-diffusive considering the spatiotemporal limitations of the experimental setup and resolution of the fitted two-dimensional Gaussian function, since most led to *R*^2^ << 0.9. The calculation of the mean *μ* and standard deviation *σ* of the diffusion constant was done using the log-transformation due to its skewed normal distribution, as

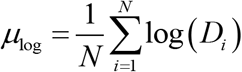

and

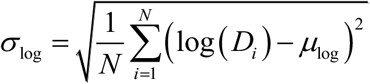

where *N* is the number of diffusion constants obtained for one experimental condition. The back-transformation was calculated then as

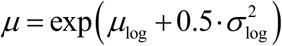

And

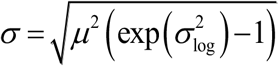

respectively, following the Finney estimator approach (Finney, 1941). The standard error of mean (*SEM*) considering a confidence interval of 95% was calculated as

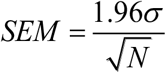

## Acknowledgments

We thank Beate Nitschke for help with protein preparation, and Stephan Eisler and Ke Zhou for help with the TIRF microscopy. We thank Robin Ghosh, for stimulating discussion, and the Baden-Württemberg Foundation for funding (BiofMO-6, SN).

## Author contributions

SN initiated and directed the study. SW fluorescently labeled proteins, collected and processed the TIRF data. MH and SW wrote the software used for data and statistical analysis. SW, MH and SN analyzed results. SN wrote the initial paper draft and secured funding. SW, MW, MH and SN edited and reviewed the draft. LF, SL and MW provided initial expertise.

## Supplementary Materials

**Fig. S1.**
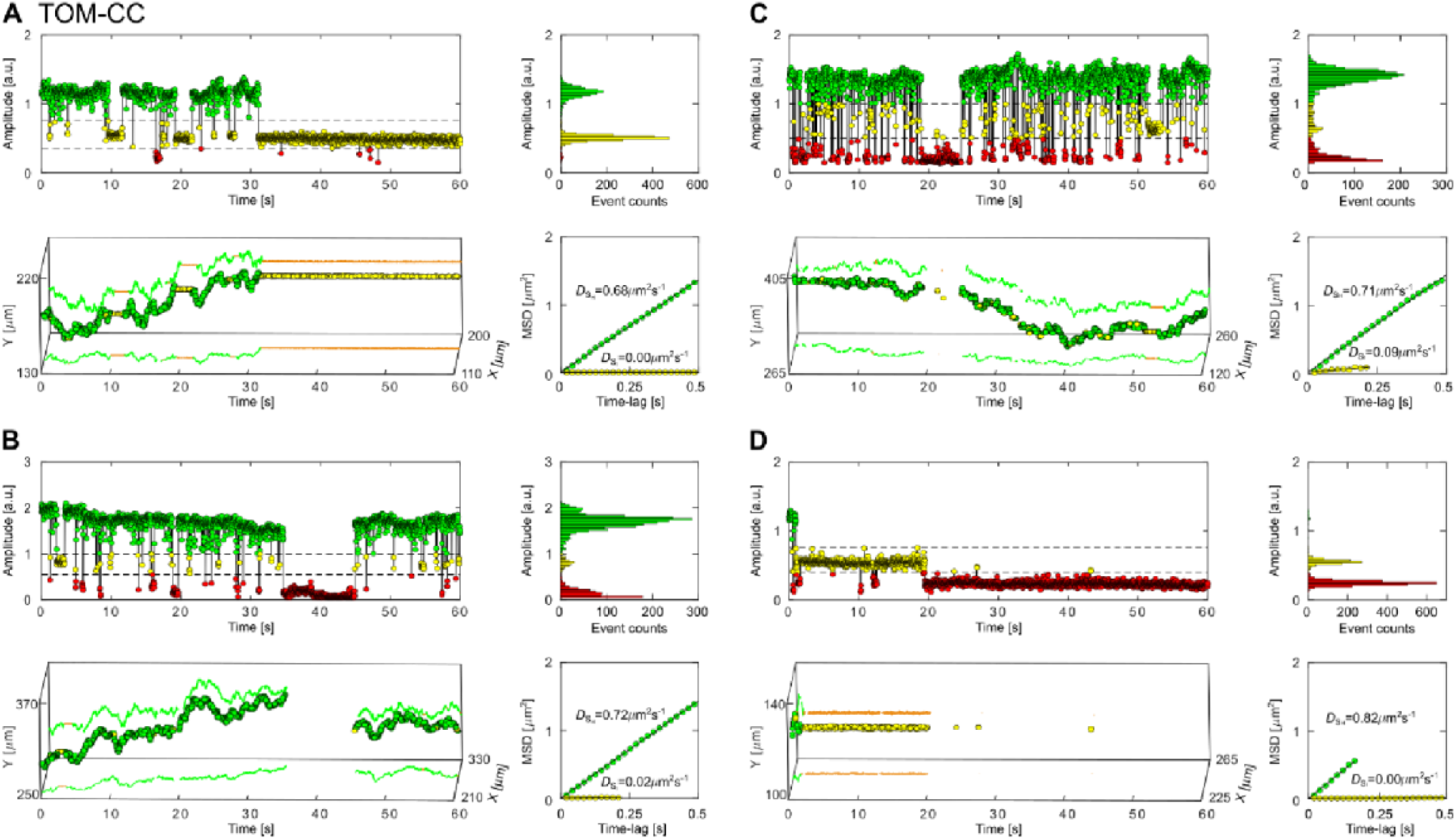
Lateral mobility correlates with the open-closed channel activity of TOM-CC. (A, B, C and D) Representative fluorescent amplitude traces of individual TOM-CC channels recorded in nonmodified agarose-supported DIB membranes. The amplitude recordings and trajectories indicate that the open-closed channel activity of TOM-CC correlates with the lateral membrane mobility of the complex. The fluorescent amplitude traces (upper left) and corresponding amplitude histograms (upper right) show three distinct ion permeation states (S_H_, green; S_I_, yellow; S_L_, red). The trajectories (bottom left) display two mobility states, moving (green) and non-moving (yellow). The mean square displacement (MSD, bottom right) increases linearly with time when TOM-CC is in S_H_ state. The MSD does not change with time for TOM in the S_I_ state. Due to limited resolution, trajectory points and MSD-plots are not shown for TOM-CC in the low permeation state S_L_.

**Fig. S2.**
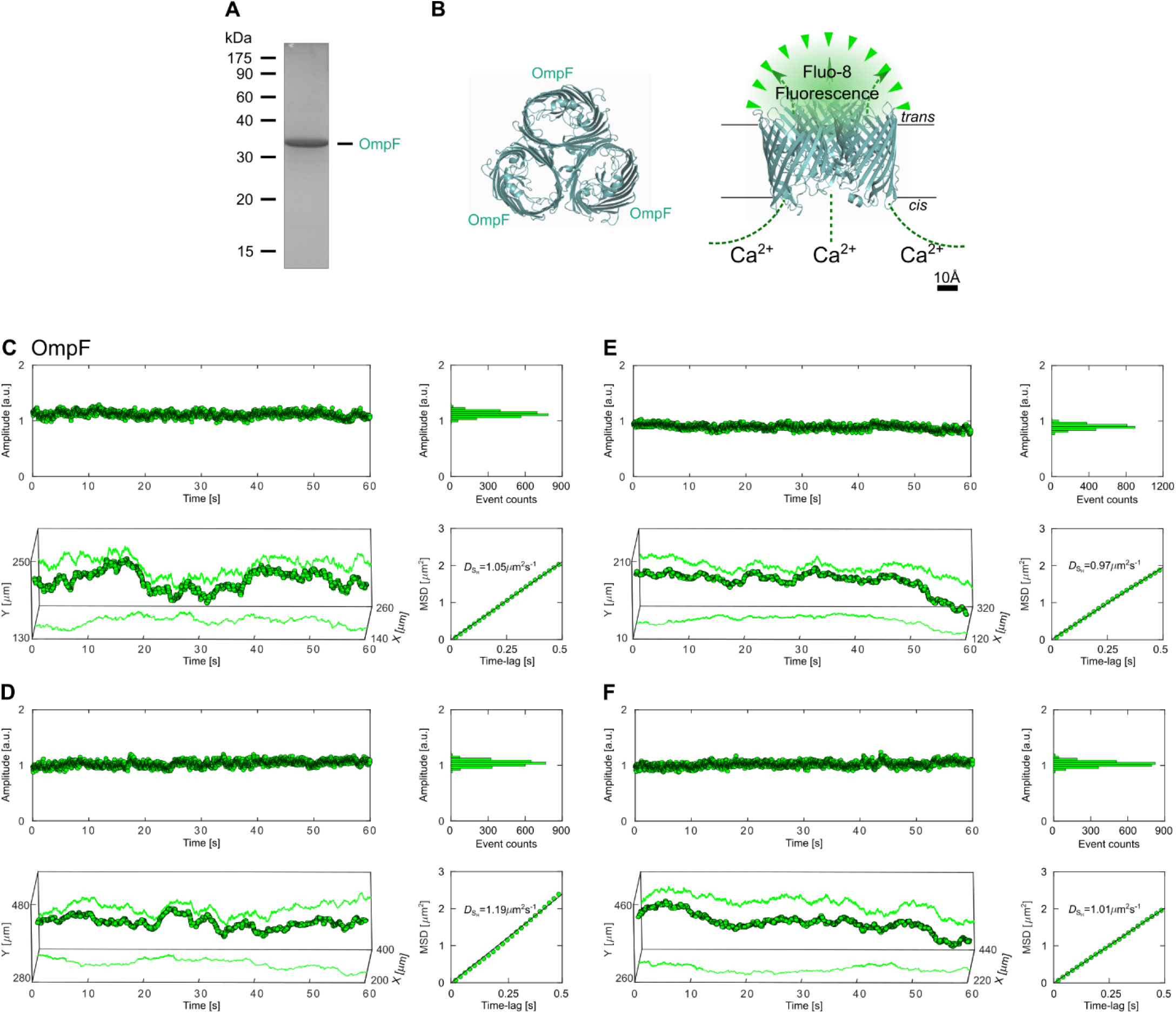
Lateral mobility does not influence the open-closed channel activity of OmpF. (A) SDS-PAGE of OmpF. The protein was purified from *E. coli* membranes by differential SDS extraction at 50°C and 37°C, respectively. (B) Atomic model of OmpF (PDB, 1OPF). Left, view from the extracellular side; right, side view. Ca^2+^ flux through the three β-barrel pores is measured by electrode-free optical single channel recording (Ef-oSCR) using Fluo-8 as Ca^2+^-sensitive dye, as described in Fig. 1. (C, D, E and F) Representative fluorescent amplitude traces of individual OmpF channels (*n* = 42) recorded in agarose-supported DIB membranes. The amplitude recordings and trajectories of the four examples indicate that the open-closed channel activity of OmpF does not correlate with the lateral mobility of the protein. OmpF displays only one ion permeation and lateral membrane mobility state, despite the fact that three channels are indicated in the atomic structure.

**Fig. S3.**
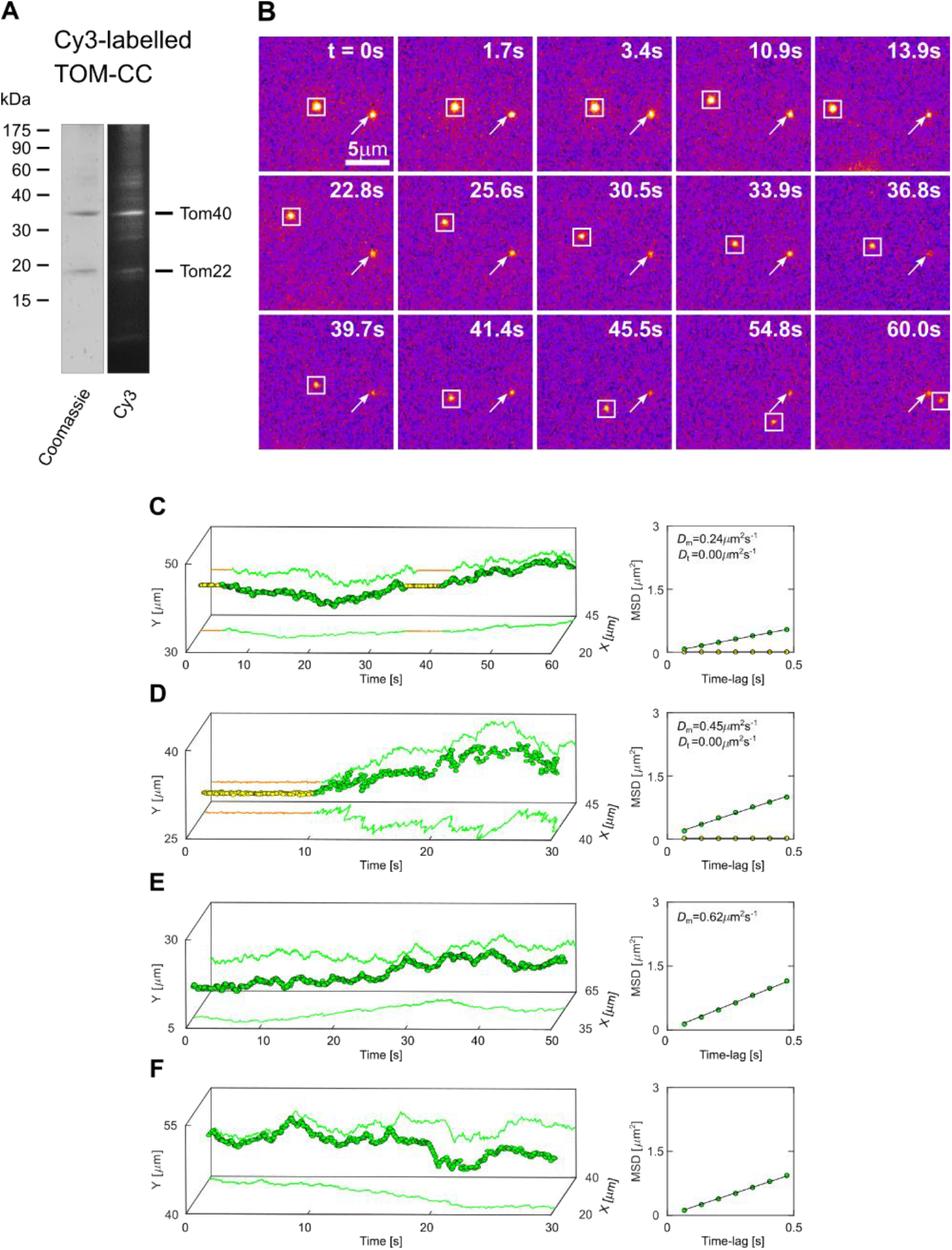
Tracking fluorescent-labeled TOM-CC in non-modified agarose supported DIB membranes. (A) SDS-PAGE of Cy3-labeled TOM-CC analyzed by Coomassie Blue staining and Cy3 fluorescence, respectively. (B) Representative TIRF microscopy images of a DIB membrane with two Cy3-labeled TOM-CC molecules (*n* = 15) taken from a time series of 60 s (Movie S4). The square-marked spot displays lateral motion, interrupted by a transient arrest between *t* = 0 s and *t* = 3.3 s, and between *t* = 33.9 s and *t* = 39.4 s. The arrow-marked spot corresponds to a non-moving TOM-CC. (C) Trajectory and diffusion coefficient of the squaremarked Cy3-labeled TOM-CC. (D, E and F) Trajectories and diffusion coefficients of additional Cy3-labeled TOM-CCs. The color-coding of trajectories is the same as in Fig. 3. Note that the freely moving TOM-CC molecule in Figures B and C stops at the same spatial x,y-position when it sweeps this position a second time, indicating a specific molecular trap or anchor point at this position below the membrane.

**Fig. S4.**
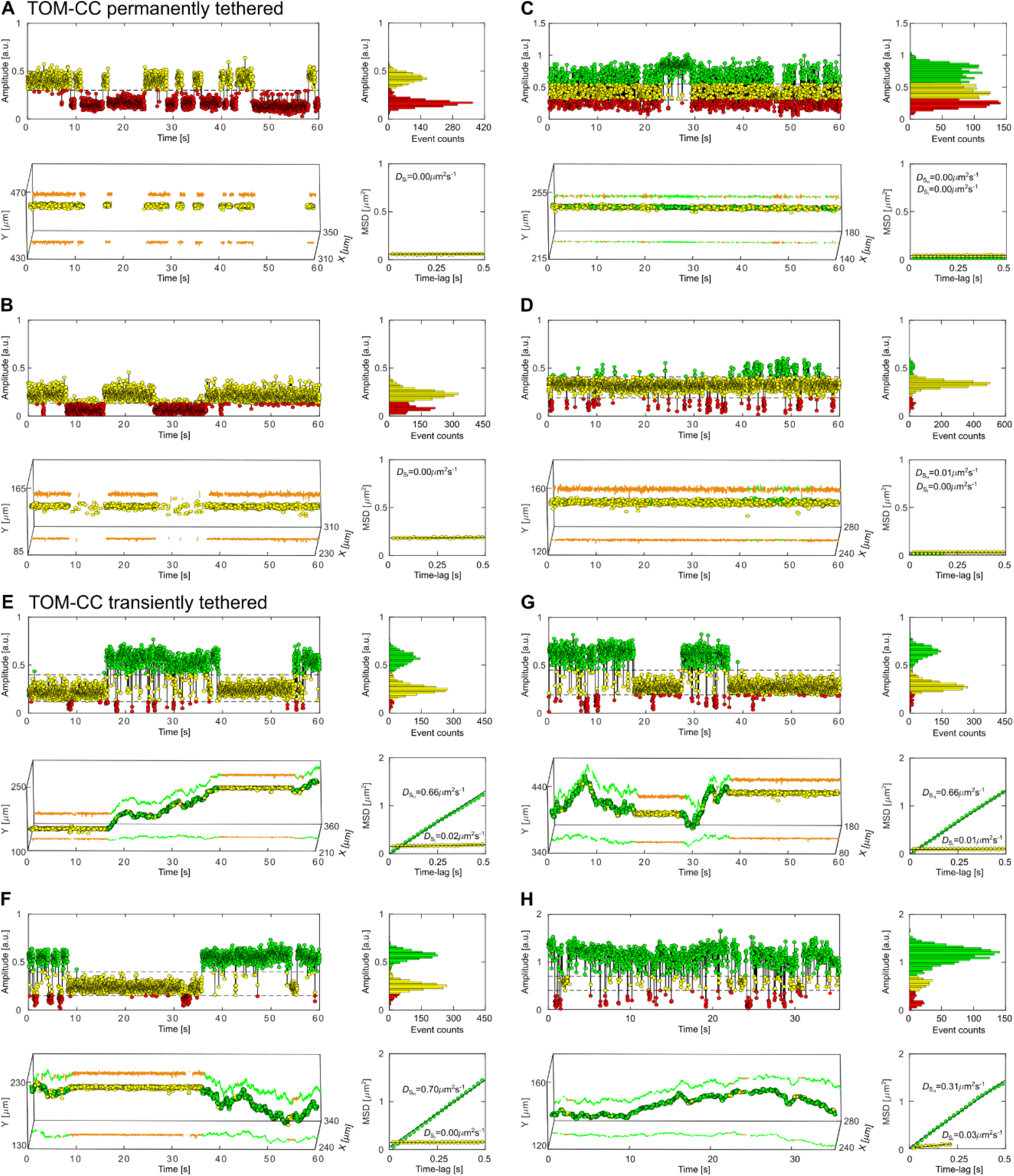
Controlled immobilization of His-tagged TOM-CC triggers channel closures. Representative fluorescent amplitude traces of individual TOM-CC channels (*n* = 109) recorded in Ni-NTA-modified agarose-supported DIB membranes. (A, B, C and D) Particles are permanently trapped. (E, F, G and H) Particles are transiently trapped. Fluorescent amplitude traces (upper left), amplitude histograms (upper right), trajectories (bottom left) and mean square displacement plots (MSD, bottom right) are similar to those shown in Figs. 4B and 4C, respectively. The color-coding is the same as in Fig. 2.

**Fig. S5.**
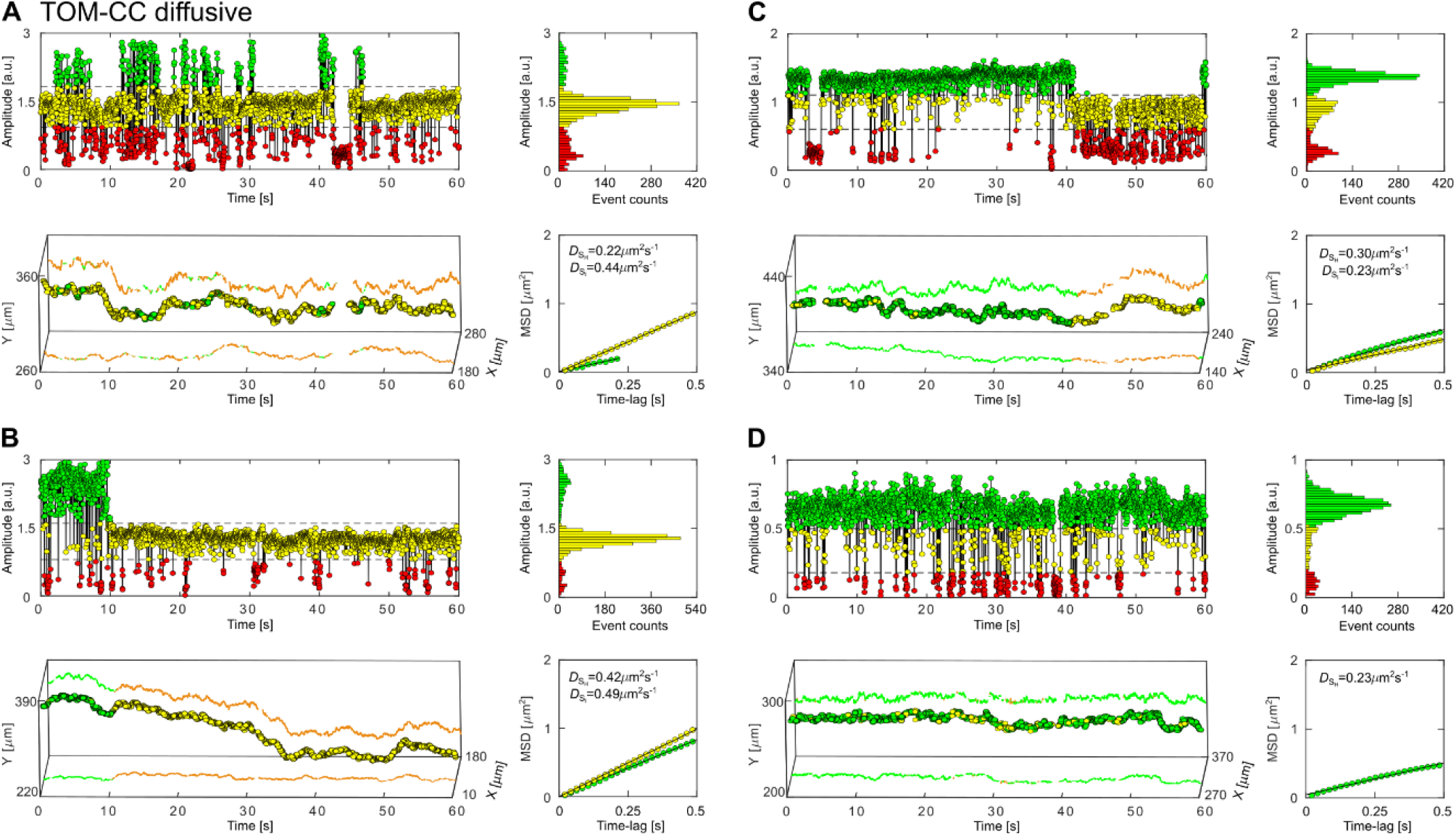
Lateral mobility and open-closed channel activity of TOM-CC in Ni-NTA-modified agarose supported DIB membranes. (A, B, C and D) Fluorescent amplitude traces (upper left), amplitude histograms (upper right), trajectories (bottom left) and mean square displacement plots (MSD, bottom right) are shown. Particles (*n* = 14) are continuously moving and represent nontypical TOM-CCs. The color-coding is the same as in Fig. 2.

**Movie S1. Imaging the channel activity of single TOM-CC molecules, related to Fig. 2A**. A Ca^2+^ indicator dye (Fluo-8) is used to monitor Ca^2+^-flux through individual TOM-CC channels in a non-modified agarose-supported DIB membrane using electrode-free optical single channel recording (Ef-oSCR). TOM-CCs appear as bright spots under 488 nm TIRF-illumination. The spots show high and intermediate intensity corresponding to two conformational states S_H_ (green) and S_I_ (yellow) with two pores and one pore open, respectively. The low intensity level represents a conformation S_L_ (red) with both pores closed. Raw image data are shown; individual spots are marked according to their conformational states.

**Movie S2. Time evolution of TOM-CC channel activity, related to Figs. 2B - 2C**. Fitting the fluorescence intensity profiles of a single TOM-CC to two-dimensional Gaussian functions (right). Red, yellow and green intensity profiles represent TOM-CC in S_L_, S_I_ and S_H_ demonstrating Tom40 channels, which are fully closed, one, and two channels open, respectively. Original fluorescence intensities (left) were recorded at a pixel size of 0.16 μm and at a frame rate of 47.51 s^−1^. No bleach correction and filter algorithm were applied.

**Movie S3. Correlation between stop-and-go dynamics and open-closed channel activity of single TOM-CC molecules, related to Fig. 3**. TIRF image recording (left) and trajectories (right) of individual TOM-CC molecules in a non-modified agarose supported DIB membrane. The square-marked spots display lateral motion (Go) interrupted by transient arrest (Stop). The red, yellow and green color coding corresponds to TOM-CC molecules, which are fully closed (S_L_), one (S_I_) and two (S_H_) channels open, respectively. Moving TOM-CC molecules in S_H_ switch to S_I_ or S_L_ when they stop in the DIB membrane. Raw image data are shown; individual spots are marked according to their conformational states (left). The trajectories of moving TOM-CC molecules are colored in green; the trajectories of trapped TOM-CC molecules in S_I_ are colored in yellow; the trajectories of trapped TOM-CC molecules in S_L_ are not shown because weak intensity profiles do not allow accurate determination of the position of TOM-CC in the membrane plane.

**Movie S4. Stop-and-go movement of fluorescently labeled TOM-CC, related to Fig. S3**. TIRF image recording (left) and trajectories (right) of Cy3-labeled TOM-CC molecules in a non-modified agarose supported DIB membrane. The square-marked spots display lateral motion (Go, green), interrupted by a transient arrest (Stop, yellow). Note that the freely moving TOMCC molecule stops at the same spatial x,y-position (yellow cross) when it crosses the same position a second time, indicating a specific molecular trap or anchor point at this position below the membrane. Raw image data are shown. Grey scales of individual images are transformed into pseudo colour images to better display movement of fluorescently labelled TOM-CC molecules. No bleach correction and filter algorithm were applied.

**Movie S5. Lateral movement and channel activity of single OmpF molecules, related to Figs. 2E - 2F, and Fig. S2**. TIRF image recording (left) and trajectories (right) of single OmpF molecules in a non-modified agarose supported DIB membrane. A Ca^2+^ indicator dye (Fluo-8) is used to monitor Ca^2+^-flux through three-pore β-barrel OmpF channels using electrode-free optical single channel recording (Ef-oSCR). OmpF appears as bright spots under 488 nm TIRF illumination. The channel activity of OmpF does not correlate with the lateral mobility of the protein. OmpF displays only one ion permeation and lateral membrane mobility state. Raw image data are shown and individual spots are marked.

**Movie S6. Single channel activity of transiently and permanently trapped TOM-CC molecules, related to Figs. 3, 4 and 5**. TIRF image recordings (left) and trajectories (right) of single TOM-CC molecules in non-modified (top) and Ni-NTA-modified (bottom) agarose supported DIB membranes. The spots show high and intermediate intensity corresponding to two conformational states S_H_ (green) and S_I_ (yellow) with two pores and one pore open, respectively. The low intensity level represents a conformation S_L_ (red) with both pores closed. Diffusive TOM-CC molecules are only in S_H_ state. Non-diffusive molecules are either in S_I_ or S_L_. The permanently tethered fraction of TOM-CC in Ni-NTA-modified agarose is significantly larger compared to the fraction in non-modified agarose over time. Raw image data are shown; individual spots are marked according to their conformational states; trajectories of trapped TOM-CC molecules in S_L_ are not shown because weak intensity profiles do not allow accurate determination of the position of TOM-CC in the membrane plane. Original fluorescence intensities were recorded at a pixel size of 0.16 μm and at a frame rate of 47.51 s^−1^.

